# Compression Efficiency and Structural Learning as a Computational Model of DLN Cognitive Stages

**DOI:** 10.64898/2026.02.01.703168

**Authors:** Alia Wu

## Abstract

We propose a computational instantiation of three cognitive stages from the Dot–Linear– Network (DLN) framework, grounded in a compression-efficiency thesis. DLN stages are characterized as graph-structured belief-dependency representations used to evaluate options: Dot as no persistent belief graph (reactive policies with negligible internal state), Linear as a null graph over option beliefs (*K* independent option estimates with no information sharing), and Network as shared latent structure (a bipartite factor graph in which *F* latent factors connect to *K* options), augmented by a temporal exposure state and an explicit structural learning cycle (hypothesis → test → update/expand).

We distinguish two compression targets—option-factor structure (shared components in expected outcomes) and stakes-factor structure (shared drivers of consequence-bearing exposures)— whose intersection yields jointly efficient actions that simultaneously improve expected outcomes and marginal exposure impact. In a bandit-like simulation (100 seeds, *K* ∈ { 20, 50, 100, 200 }, *F* =5), Network policies dominate Linear policies in cost-adjusted utility at large *K*, with the empirical crossover occurring much earlier than an analytic cost-only prediction (*K*^∗^ = *F* + *c*_meta_*/c*_param_), revealing that the advantage is primarily statistical (shrinkage-like estimation gains from factor pooling) rather than purely computational. Under stakes, all non-DLN agents—including Linear-Plus agents with identical factor structure and Network-standard agents with hierarchical Bayesian learning—collapse due to unmodeled cumulative exposure, while Network-DLN maintains positive utility. Within-stage consistency tests (two algorithmically distinct agents per stage) confirm that the collapse pattern is determined by representational topology, not algorithmic choice.

These results evaluate internal consistency of a DLN-to-computation mapping under explicit assumptions; they do not validate a developmental theory in humans.

## 1 Introduction

Individuals can possess comparable domain knowledge yet differ sharply in *how* that knowledge is organized. One person may represent a situation as a set of disconnected facts; another as an interacting system with feedback and cross-constraints—a distinction central to systems thinking [38, 30]. These differences matter: two individuals with equivalent factual knowledge can make systematically different choices because their representations differ in what they treat as shared, what they treat as independent, and how constraints propagate across elements. Making such structural differences explicit is relevant to human–AI interaction and cognitive modeling [27].

We propose the Dot–Linear–Network (DLN) framework, which treats such differences as differences in *representational topology*. A cognitive state is represented as a graph *G* = (*V, E*), where nodes are representational elements (concepts, features, action templates) and edges are statistical, causal, or constraint relations that allow information to propagate. Our topological claims concern the agent’s *belief-dependency structure* used to evaluate options. In this framework, Dot corresponds to no persistent belief structure (reactive baselines), Linear corresponds to a null graph over *K* option beliefs (independent option-by-option estimates with no edges), and Network corresponds to shared latent structure (options are d-connected through a smaller set of *F* latent factors). Temporal coupling through cumulative exposure and a meta-level revision loop create feedback at the level of *dynamics and learning*, even when the within-timestep belief graph is acyclic. This complements work on knowledge graphs [19], semantic networks [8, 1], and cognitive architectures [2, 26] by foregrounding the dependency structure of what agents know, not just the content.

This paper focuses on a specific computational thesis: **Network representations are more efficient in stable environments because they represent shared components once rather than redundantly across each branch**. Exploration strategies are well-studied in bandits and reinforcement learning [13, 37, 42]; our focus is the representational *reuse* mechanism that can render Network efficient even in stable environments.

We decompose the thesis into two compression targets. First, *option-factor structure*: many options share components in their expected outcomes, so learning can be amortized across options if shared structure is represented (hierarchical shrinkage provides a classical example [20, 9]). Second, *stakes-factor structure*: the consequences of actions often share common drivers (exposure factors) in addition to idiosyncratic effects. When consequences are coupled through shared drivers, marginal impact depends on cumulative exposure, so a decision policy must represent cross-terms (covariances) rather than treating each action’s stakes in isolation. The intersection of these structures makes *jointly efficient actions* possible: a single action can be simultaneously beneficial for expected outcomes and beneficial for marginal stakes impact.

We operationalize Network as a factorized model class coupled to a structural learning cycle (detailed in Section 5.1). When the structural hypothesis is correct, the agent exploits compression and achieves *O*(*F*)-like scaling in the number of shared factors *F* ; when the hypothesis fails, the agent pays additional cost to recover by expanding toward *O*(*K*) state.

### Contributions

This paper contributes: (1) an explicit DLN-to-computation mapping centered on factorized compression rather than entropy-based exploration; (2) a stylized environment with option-factor structure, stakes-factor structure, and an explicit cross-term that makes cumulative exposure matter; (3) scaling results with paired-seed statistical summaries (bootstrap confidence intervals and paired effect sizes) identifying where compression helps and where it does not; and (4) an explicit assumption audit and reproducible code.

The simulation evaluates internal consistency of this mapping; it does not establish that human cognitive development follows DLN.

### Paper organization

Section 2 formalizes DLN stages as topological regimes and introduces the model space and revision graph. Section 3 states claims, diagnostic signatures, and testable behavioral predictions. Section 4 situates the work relative to existing approaches. Section 5 describes the simulation design. Section 6 presents results, including within-stage consistency tests across DLN stages and the contraction mechanism. Section 7 discusses interpretation, boundary conditions, and limitations. Section 8 describes reproducibility procedures. Section 9 concludes.

## 2 DLN as a topological framework

We model a cognitive state as a graph *G* = (*V, E*) where nodes are representational elements (features, concepts, action templates) and edges are relations (associations, causal links, constraints).

### 2.1 Topological regimes

DLN describes a progression from Dot to Linear to Network, with increasingly connected representational graphs. In this paper, the relevant object is the agent’s *belief-dependency structure* used to evaluate options. The Linear stage corresponds to independent option-by-option estimation.

#### Dot stage (no persistent belief graph)

Dot corresponds to policies with no persistent belief state for option evaluation. In our simulation, Dot selects uniformly at random and does not maintain outcome or exposure estimates. Topologically, this corresponds to an empty (or ephemeral) belief graph: no stored nodes or edges allow information to propagate across options.

#### Linear stage (independent option beliefs; null belief graph)

Linear corresponds to maintaining *K* independent option beliefs (per-option means) with no edges between them. Topologically, this is a null graph on *K* belief nodes: no option shares an ancestor with any other option, so learning about option *i* does not update beliefs about option *j*. This captures the DLN property of branch-by-branch processing without reuse.

#### Network stage (shared latent structure; bipartite factor DAG)

Network corresponds to representing options as arising from a smaller set of shared latent factors. In our instantiation, the within-timestep belief graph is a bipartite directed acyclic graph (DAG): *F* latent factor nodes connect to *K* option nodes. Options that share a factor share an ancestor and are therefore d-connected through the factor; evidence from one option updates the factor belief, which updates predicted values for all factor-mates. This shared-parent structure is the graph-theoretic source of integration in our implementation: shared structure represented once and reused across many options.

#### Where cycles enter: dynamics and meta-learning

Although the factor belief graph is a DAG, Network-stage cognition in DLN also involves feedback and revisitation. In this paper, those notions correspond to (i) a closed-loop dependency in the *system dynamics* when stakes are active (actions update exposure *E*_*t*_, and *E*_*t*_ feeds back into the marginal penalty term used to evaluate future actions), and (ii) a *process-level* structural learning cycle (Section 5.1) that revises which belief graph is being used.

Section 2.4 formalizes this distinction by defining a model space ℳ whose elements are candidate belief structures and a revision graph ℛ whose edges are transitions between them; the cycle that defines Network cognition is a structural feature of ℛ, not of any individual belief graph *G* ∈ ℳ.

Investing in shared latent structure reduces the number of distinct quantities that must be learned from *K* to *F* when the structure hypothesis is correct.

### 2.2 Cost-scaling interpretation

The computational claim tested here is a scaling distinction:

- **Dot:** no accumulation of learning across encounters (reactive; baseline).
- **Linear:** independent estimation per option, yielding complexity that scales with the number of options *K*.
- **Network:** factorized representation that learns a smaller number of shared factors *F* and propagates factor updates to many options, yielding complexity that scales primarily with *F* rather than *K*.

#### Formal complexity statement

Let Linear maintain per-option sufficient statistics 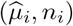 for *i* ∈ {1, …, *K*}. Then its representational state requires

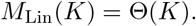

In our implementation, Linear also scores all options each step, yielding per-decision computation *G*_Lin_(*K*) = Θ(*K*). Let Network maintain per-factor statistics 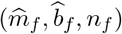 for *f* ∈ {1, …, *F*} plus the exposure state *E*_*t*_, and use a bounded within-factor local search of size *L* (treated as a constant). Then

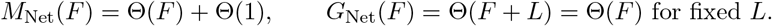

When the factor hypothesis fails and the model expands to a tabular representation, Network reverts to Θ(*K*) state and computation. Thus the compression thesis predicts an advantage only when *F* ≪ *K and* the factor hypothesis remains empirically adequate; when *K* = *F* (no shared components), the advantage should vanish, constituting a falsification criterion for the compression thesis.

The cost-scaling analysis yields three testable properties that formalize when Network compression is beneficial, when it fails catastrophically, and how an agent can recover compressed operation after model failure. We state these as a single proposition whose parts are tested independently in the simulation.

#### Proposition 1

(Cost separation and recovery bound). *Let an agent face a K-option decision environment with F-factor structure (F* ≤ *K). Let c*_param_ *be the per-element parameter learning cost and c*_meta_ *be the Level 2 model-revision overhead (navigation of the revision graph R, Section 2.4)*. *Then:*

#### (i) Crossover

*For K < K*^∗^ = *F* + *c*_meta_*/c*_param_, *Linear achieves lower total cognitive cost than Network (Network cannot do better than operating in a correct compressed model and still paying its Level 2 overhead). For K > K*^∗^, *Network under a correct factor hypothesis achieves lower total cost by replacing O*(*K*) *per-episode parameter maintenance with O*(*F*), *up to the fixed overhead c*_meta_.

#### (ii) Catastrophic failure under cross-terms

*When the environment contains stakes-factor structure with covariance coupling, Linear’s utility degrades without bound in cumulative exposure E*_*t*_ *because it cannot represent the marginal penalty term* 2*E*_*t*_*b*_*i*_. *Network’s utility remains bounded because it tracks E*_*t*_ *and computes marginal impact*.

#### (iii) Recovery bound

*When the factor hypothesis is incorrect and the agent expands to G*_tab_, *an agent whose revision graph* ℛ *includes a return transition* (*G*_tab_, *G*_*F*_) ∈ 𝒯 *recovers O*(*F*) *cost scaling within n*_contract_ + *w* + *n*_converge_ *steps after factor structure becomes present, where n*_converge_ *is the number of observations required for within-factor variance to fall below the contraction threshold. An agent without a return transition (expand-only Network or Linear) remains at O*(*K*) *cost indefinitely*.

*Parts (i) and (ii) are tested by the existing simulation results. Part (iii) is tested by the contraction ablation (Section 6.8)*.

#### Argument for (i)

In the absence of structural sharing, Linear must maintain *K* parameters, so its per-episode cost is proportional to *c*_param_*K*. A Network agent that is operating in a correct compressed model maintains only *F* parameters, but still pays a fixed Level 2 overhead *c*_meta_ for monitoring and revision. Comparing *c*_param_*K* against *c*_param_*F* + *c*_meta_ yields the crossover condition *K*^∗^ = *F* + *c*_meta_*/c*_param_.

#### Argument for (ii)

Under stakes, the true marginal impact of choosing option *i* contains a cross-term proportional to 2*E*_*t*_*b*_*i*_, where *E*_*t*_ is cumulative exposure. A Linear agent that treats options independently can include a per-option penalty (e.g.,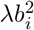) but cannot represent the coupling to *E*_*t*_, so it cannot stabilize exposure as |*E*_*t*_| grows. A Network agent that tracks *E*_*t*_ and computes marginal impact can keep exposure bounded, preventing unbounded penalty growth.

#### Argument for (iii)

After expanding to *G*_tab_, an agent with a return transition can periodically test whether a compressed factor model predicts well again (for example via a length-*w* evaluation window after a holdout period *n*_contract_). Once factor structure is present and sufficient data are accumulated (*n*_converge_ observations), the agent contracts back to *G*_*F*_ and returns to *O*(*F*) scaling. Without a return transition, the agent remains in the expanded model and continues to incur *O*(*K*) cost.

Together, parts (i)–(iii) delineate the lifecycle of a compression-based agent: initial advantage through shared structure, vulnerability when that structure fails, and—when the revision graph includes a return transition—bounded recovery. The simulation tests each part: the crossover is visible in the scaling results (Section 6.1), catastrophic failure appears in the stakes condition (Section 6.2), and recovery is tested by the contraction ablation (Section 6.8).

This relates to classical results on shrinkage and hierarchical estimation: when structure exists, sharing statistical strength across related items improves efficiency [20, 9]. In our mapping, Network operationalizes this by tracking factor-level beliefs and using them to score options, while retaining the capacity to expand to option-specific state when structure is violated.

### 2.3 Factor structure as a computational analogue of network integration

The computational instantiation represents Network as a shared factor space, rendering the topology-to-computation mapping concrete. If we represent each option *i* as connected to a factor node *c*_*i*_, then the representational graph becomes a bipartite structure in which many options share the same factor node. Updating the factor belief propagates immediately to all connected options. This is precisely the “represent shared components once” property that DLN attributes to integrated (network-stage) organization.

Stakes-factor structure adds an additional coupling: the current exposure state *E*_*t*_ links sequential decisions, and the marginal stakes impact contains a cross-term 2*E*_*t*_*b*_*i*_. This introduces a feedback dependency across time: the consequences of an action depend on what has been selected before. A Linear representation that evaluates stakes per option without tracking *E*_*t*_ treats this dependency as if it were absent; a Network representation explicitly represents the coupling and therefore computes marginal impact. In this minimal setting, the coupling induces closed-loop decision dynamics: the state induced by prior actions feeds back into the evaluation of future actions.

This mapping implies a design requirement: to test DLN Network as integrated structure, the environment must contain shared components and cross-constraints that make reuse and coupling beneficial. If structure is absent or if *K* = *F* (no shared components), the representation should not help. Those boundary tests are part of the falsifiable surface of the thesis.

### 2.4 Model space and revision graph

The preceding sections characterize DLN stages by the within-episode belief-dependency graph *G* that an agent uses to represent outcomes, exposures, and option relations. Our Network instantiation also implements an explicit structural learning cycle that can *revise which belief graph is active* when a structural hypothesis fails. To formalize this two-level distinction, we separate (i) inference and learning *within* a belief graph (Level 1) from (ii) monitoring and revising the belief graph itself (Level 2).

#### Model space

Let ℳ be a set of candidate belief-dependency structures. In the present instantiation, the core model space contains two elements: a compressed factor graph *G*_*F*_ (shared latent factors connecting options) and an expanded tabular graph *G*_tab_ (independent option beliefs). Formally, ℳ = {*G*_*F*_, *G*_tab_}.

#### Revision graph

Define the *revision graph* as a directed graph over model space, ℛ = (ℳ, 𝒯), where 𝒯 ⊆ ℳ × ℳ is the set of revision transitions available to an agent. In our simulation, the key transitions are: (i) *expansion* (switching from *G*_*F*_ to *G*_tab_ when the factor hypothesis fails) and *contraction* (switching from *G*_tab_ back to *G*_*F*_ when predictive evidence supports compression again).

#### Completing the cycle: contraction

Expansion alone yields an irreversible path *G*_*F*_ → *G*_tab_. Adding a contraction transition (*G*_tab_, *G*_*F*_) ∈ 𝒯 completes a revision cycle. This return transition matters even when |ℳ| = 2: it is the mechanism that enables an agent to recover compressed *O*(*F*) operation after a period of operating in an expanded *O*(*K*) representation.

#### Revision capacity by DLN stage

The revision graph characterizes each stage’s meta-cognitive capacity in graph-combinatorial terms:

- **Dot**. No model space (|ℳ| = 0); ℛ is empty. No revision capacity.
- **Linear**. ℳ = {*G*_tab_}; a single vertex with no edges. The agent occupies a fixed point in model space.
- **Network (expand-only)**. ℳ = {*G*_*F*_, *G*_tab_}; one directed edge *G*_*F*_ → *G*_tab_. The agent can detect model failure and expand, but cannot return to a compressed representation.
- **Network (full cycle)**. Two directed edges (expand and contract). ℛ contains a cycle: the agent can depart from a model, operate under an alternative, and return. This is the graph-theoretic signature of full revision capacity.

The functional consequence of the return transition is stated in Proposition 1(iii): an agent that can return to *G*_*F*_ recovers *O*(*F*) cost scaling in bounded time; an agent that cannot is permanently locked at *O*(*K*).

#### Two levels of computation

Level 1 computation concerns inference and parameter learning within a chosen belief graph *G* ∈ ℳ (e.g., updating factor means or option means). Level 2 computation concerns monitoring model adequacy and executing transitions in ℛ. Proposition 1(iii) establishes that an agent whose revision graph includes a return transition recovers Θ(*F*) scaling in bounded time once factor structure is present, while an agent without a return transition remains at Θ(*K*) permanently.

## 3 Claims and predictions

### 3.1 Claims and diagnostic signatures

The computational mapping yields diagnostic signatures that are testable in simulations (as demonstrated here) and in behavioral experiments (Section 3.2):

- **C1: Scaling signature (option-factor compression)**. In tasks with option-factor structure, cost-adjusted utility should scale with *F* under Network representations but with *K* under Linear representations, because Network learns shared structure once and reuses it across options.
- **C2: Marginal-stakes signature (stakes-factor compression)**. When consequences depend on cumulative exposure (covariance cross-terms), Network policies should evaluate *marginal* impact given current exposure state *E*_*t*_, while Linear policies evaluate each option’s consequences independently and miss covariance-driven marginal effects.
- **C3: Intersection signature (jointly efficient actions)**. When option-factor and stakes-factor structure overlap, Network policies should identify actions that serve both objectives simultaneously, because both decompositions are represented in a shared factor space.
- **Prior failure signature (recovery)**. Under prior-wrong conditions, Network policies should exhibit measurable model updates (switches/expansions) and improved fit relative to no-test or no-update ablations, at the cost of increased cognitive cost.

These signatures separate the compression thesis from generic claims of adaptation or exploration. The scaling signature should obtain in stable environments without distribution shift.

### 3.2 Testable behavioral predictions

The diagnostic signatures above yield concrete predictions for human behavioral experiments. We state three that are directly testable with existing paradigms and require no new simulation results.

#### Response-time scaling

In a multi-option choice task with labeled factor categories (e.g., product sectors) and varying *K* (holding *F* fixed), the model predicts that participants using Network-stage representations should exhibit response times that are approximately flat in *K*, because evaluation proceeds at the factor level (*O*(*F*) comparisons) with bounded local search within the selected factor. Participants using Linear-stage representations should exhibit response times that scale with *K*, because each option is evaluated independently. A within-subject design in which the same participant encounters option sets of varying size would allow estimation of the scaling exponent, with the DLN prediction being a sharp qualitative contrast: sublinear vs. linear growth. Process-tracing paradigms (e.g., Mouselab or eye-tracking) could further distinguish factor-level scanning patterns from exhaustive option-level scanning.

#### Transfer and generalization

After training on factor-structured options, introduce novel options that share a known factor label but have not been individually encountered. A Network-stage participant should generalize at *O*(*F*) rather than *O*(*K*) cost: the factor-level estimate transfers to the new option without requiring item-specific learning, yielding above-chance performance from the first trial. A Linear-stage participant should show no such transfer, because the novel option has no associated option-level estimate. This prediction distinguishes factor-based generalization from item-specific learning and can be tested in standard categorization-and-choice paradigms.

#### Cumulative-exposure sensitivity

Under tasks where cumulative exposure produces real or simulated consequences (e.g., portfolio allocation with cumulative risk), the model predicts that Network-stage decision-makers should reverse option preferences when cumulative exposure *E*_*t*_ crosses sign thresholds—selecting options with *b*_*i*_ *<* 0 when *E*_*t*_ *>* 0, and vice versa—because they evaluate the marginal cross-term 2*E*_*t*_*b*_*i*_. Linear-stage decision-makers should exhibit fixed preference patterns that are independent of *E*_*t*_, because their scoring function 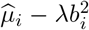 does not reference cumulative state. This yields a testable interaction: exposure history should modulate option choice for Network-stage but not Linear-stage participants.

### 3.3 Explicit assumptions

We state assumptions explicitly, as they determine the scope of claims the simulation can support.

- **A1 (observed option-factor labels)**. Each option *i* has an observed option-factor label *c*_*i*_ ∈ {0, …, *F* − 1}. This stands in for a shared-component representation. (Linear agents choose to ignore it.)
- **A2 (observed stakes loading)**. Each option exposes an observed stakes loading *b*_*i*_ ∈ ℝ.
- **A3 (structured outcomes; prior correct)**. In the structured condition, mean outcomes follow a factor model: 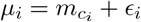.
- **A4 (unstructured outcomes; prior wrong)**. In the unstructured condition, *µ*_*i*_ are independent and the label *c*_*i*_ is irrelevant for predicting outcome.
- **A5 (stakes objective and covariance cross-term)**. Stakes are represented as a quadratic exposure objective in a 1D exposure state *E*_*t*_. Selecting option *i* updates exposure *E*_*t*+1_ = *E*_*t*_ + *b*_*i*_ and incurs incremental penalty

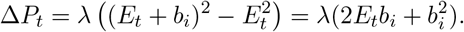

The cross-term 2*E*_*t*_*b*_*i*_ is the minimal mechanism by which cumulative exposure creates marginal effects that cannot be captured by option-by-option penalty terms alone.

- **A6 (cognitive cost proxy)**. We use an algorithmic proxy cost consisting of memory units and compute units with weights (reported in the run manifest). This is a complexity surrogate, not a claim about biological energy.

## 4 Relationship to existing work

### 4.1 Cognitive architectures and bounded rationality

DLN provides a structural lens that complements established cognitive theories and architectures. Many cognitive architectures specify symbolic or production-system mechanisms [2, 26], and many modern AI systems operationalize relational knowledge as graphs [19]. The present contribution emphasizes that belief-dependency structure imposes computational constraints. When option beliefs are independent, evidence updates a single option; when options share latent structure, evidence propagates through shared parents, allowing one observation to update many options. This constitutes a representational and statistical-generalization claim rather than a claim about the computational cost of cyclic belief graphs.

DLN also connects to bounded rationality traditions in which limited computational resources necessitate simplified representations [39, 48, 15]. In this paper, the resource constraint is made explicit as a cognitive cost proxy and a scaling claim: Linear representations are expensive because they duplicate estimation across options, while Network representations can amortize learning across options when shared structure exists.

From a developmental perspective, stage theories often describe qualitative shifts in what operations become possible [34, 21]. The distinctive DLN claim concerns a shift in representational topology that makes certain computations cheaper (reuse) and others unnecessary (redundant branch-by-branch processing). Dynamic-systems accounts emphasize that qualitative shifts can emerge from continuous changes in coupling strength and feedback [45]. The structural learning cycle is compatible with this view: representational coupling is strengthened when it explains data and relaxed when it fails.

### 4.2 Computational neuroscience: structure learning as a distinct operation

Computational neuroscience and cognitive science have repeatedly distinguished structure learning (selecting or revising a latent representation) from parameter learning (updating values within a chosen representation) [14, 7, 22, 12, 24]. Neural evidence supports this distinction: structure learning signals in frontoparietal cortex are at least partially dissociable from striatal value-learning signals [47, 28, 23].

Building on this foundation, we instantiate DLN stage differences as differences in *belief-dependency structure* and derive concrete efficiency predictions that are tested against seven baseline agents drawn from established computational traditions, including Thompson sampling [37], Rescorla– Wagner associative learning [35], hierarchical Bayesian shrinkage [14], and online latent-factor inference [14]. First, we make the scaling implication of shared latent structure explicit as an *O*(*F*) versus *O*(*K*) crossover (Section 2.2), where *F* is the number of shared factors and *K* is the number of options. Second, we extend the same compression logic to *stakes-factor structure* by introducing an exposure objective with an explicit covariance cross-term, so that the marginal consequence of an action depends on cumulative exposure state. Third, we connect these representational regimes to a three-stage DLN mapping (Dot/Linear/Network) and test falsifiers and ablations (e.g., *K*=*F* and prior-wrong conditions) that isolate when the structural learning cycle matters.

This positioning also clarifies how our claims relate to graph-theoretic complexity results. Classical work on probabilistic graphical models [33] and the treewidth / tractable-cognition literature [49, 25] analyze the complexity of inference *within a fixed model structure* and establish that low-treewidth graphs enable efficient inference. The within-timestep belief graph for the Network agent is itself a low-treewidth bipartite DAG. Our claim is a representation-construction question: when the world admits shared structure with *F* ≪ *K* and the factor hypothesis is empirically adequate, representing shared structure reduces the number of distinct quantities that must be learned from *K* to *F* ; when structure is absent or incorrect, expansion toward a tabular representation removes the advantage.

#### Formalizing the gap

The structure-learning literature distinguishes learning *within* a chosen representation from learning *which* representation to use (or when to switch) [22, 7]. Section 2.4 makes this distinction explicit in our DLN instantiation: Level 1 computation occurs within a belief-dependency graph *G* ∈ ℳ, while Level 2 computation is navigation in the revision graph ℛ over model space. The crossover *K*^∗^ = *F* + *c*_meta_*/c*_param_ (Proposition 1(i)) makes precise when paying a fixed Level 2 overhead is cost-effective relative to the *O*(*F*) savings from representing shared structure.

#### Distinction from standard model selection

The present framework contributes three properties absent from standard model selection. First, revision is *costly* : expansion and contraction incur explicit cognitive costs (switch cost *c*_switch_, memory reallocation) that enter the agent’s objective, making representation change an economic tradeoff rather than a free comparison. Second, revision capacity is a property of the *agent*, not a universal procedure: Linear agents face the same model space ℳ but lack revision transitions (|𝒯 | = 0), and the ablation results (Section 6) demonstrate that this constraint produces measurably different performance. Third, the revision graph ℛ is itself a cognitive variable whose topology—empty, acyclic, or cyclic—varies across agents, rather than a fixed algorithmic step. By contrast, standard Bayesian model comparison (e.g., via marginal likelihood or BIC) treats model switching as computationally free, applies a universal comparison procedure, and does not represent the structure of transitions between models. The Linear-Plus agents (Thompson-Factor, RW-Feature; Section 6) demonstrate this gap empirically: agents with identical factor structure but no revision cycle or exposure tracking collapse under stakes, confirming that the properties above contribute beyond what standard factor-level learning provides.

### 4.3 DLN as a classification axis for existing models

The models classified in Table 3 span multiple literatures that rarely cite one another. Thompson Sampling researchers do not typically reference causal discovery work. Bandit researchers do not cite world model papers. Meta-learning and hierarchical Bayes address similar structural problems but appear in different venues with different formalisms. Each community has developed models that, under the DLN lens, occupy the same topological stage for the same structural reason: independent option-level estimation (Linear), designer-specified cross-option structure (Linear-Plus), or agent-discovered shared latent structure (Network). Yet no prior work has organized them along a single representational-topology axis. The DLN framework provides this axis.

**Table 1:** Assumption audit: what is built in, and relaxation paths.

| ID | Assumption (informal) | Built-in? | Relaxation path |
| --- | --- | --- | --- |
| A1 | Factor label $c_i$ is observed (shared-component index). | Yes | Learn $c_i$ / latent factors via clustering or representation learning. |
| A2 | Stakes loading $b_i$ is observed. | Yes | Learn $b_i$ (or exposures) from observed consequences. |
| A3 | Structured outcomes follow $\mu_i = m_{c_i} + \epsilon_i$. | Swept | Sweep structure strength (within-factor variance). |
| A4 | Unstructured outcomes violate the factor prior (prior wrong). | Swept | Add partial structure and drifting structure regimes. |
| A5 | Stakes penalty contains cross-term via exposure state $E_t$. | Yes | Move to multi-dimensional $E_t$ and general quadratic forms. |
| A6 | Cognitive cost is a proxy based on memory/compute scaling. | Yes | Sensitivity analysis; report Pareto frontiers instead of a single $\alpha$. |

**Table 2:** Claim-to-test matrix: what each experiment condition identifies.

| Claim | Condition | Observable signature |
| --- | --- | --- |
| C1 (Option-factor compression) | Structured, stakes= 0, sweep $K$ | Network surpasses Linear at large $K$ in cost-adjusted utility; Linear cost grows with $K$ while Network cost is $\approx O(F)$. |
| C2 (Stakes-factor compression) | Structured, stakes> 0, sweep $K$ | Linear utility degrades substantially due to unmodeled exposure cross-terms; Network remains stable by tracking cumulative exposure and using marginal impact. |
| C3 (Intersection) | Structured, stakes> 0 | Network selects jointly efficient actions (high outcome, favorable marginal stakes) at a higher rate than Linear, especially when overlap is sparse. |
| Recovery (prior wrong) | Unstructured, stakes= 0 (and > 0) | Network-Full shows switch/expansion events and outperforms NoTest/NoUpdate ablations, paying increased cognitive cost. |
| Boundary (no compression) | $K = F$ | Network advantage disappears (falsifier for compression thesis). |

**Table 3:** Classification of existing computational models by DLN representational topology. Models marked *†* are simulated in this paper; all others are classified but not simulated.

| DLN Stage | Defining Property | Computational Exemplars | Key References |
| --- | --- | --- | --- |
| **Dot** — No persistent belief graph | No belief state persists across encounters | Random policy / ε-greedy (no memory); Win-Stay Lose-Shift; Reactive/reflex agents; Satisficing heuristics | [32]; [36]; [39] |
| **Linear** — $K$ independent estimates, null belief graph | Per-option statistics, no cross-option sharing | Thompson Sampling; UCB1<sup>†</sup>; EXP3; Tabular Q-Learning; Naive Bayes; A/B Testing (frequentist) | [46]; [3]; [4]; [50] |
| **Linear-Plus** — Designer-specified cross-option structure | Fixed feature/factor representation supplied by designer; no structural learning | Correlated Thompson Sampling; GP-UCB / GP bandits; Contextual bandits (LinUCB); Deep Q-Networks; Thompson-Factor<sup>†</sup>; RW-Feature<sup>†</sup> | [41]; [29]; [31]; [35] |
| **Network** — Shared latent structure with structural learning | Agent-discovered or given hierarchy with cross-option information propagation | Hierarchical Bayesian models; HierBayes<sup>†</sup>; LatentFactorEM<sup>†</sup>; Bayesian structure learning; Causal discovery (PC, GES); MAML / Meta-learning; World models; Dirichlet Process Mixtures; Neural Architecture Search; Network-Full<sup>†</sup> | [14]; [44]; [17]; [40]; [6]; [10]; [16]; [43]; [51] |

**Note.** Linear-Plus is not a formal DLN stage but an empirically useful category: models that share information through designer-specified structure without agent-driven structural learning. Thompson-Factor and RW-Feature (this paper) are placed in Linear-Plus; HierBayes and LatentFactorEM are placed in Network (standard); Network-Full is the only agent implementing all three DLN Network mechanisms: factor compression, cumulative exposure tracking, and the structural learning cycle.

We identify an empirically important intermediate category: models that introduce cross-option information through designer-specified structure rather than agent-discovered structure. GP-UCB [41], for example, shares information across arms via a kernel function, but the kernel is chosen by the designer, not learned from interaction. The agent performs parameter updates within a fixed correlation structure. Contextual bandits [29] generalize across arms through a given feature representation, but cannot discover features the designer did not provide. Under DLN, these constitute Linear cognition with enriched parameterization, not Network cognition, because the representational topology itself is never revised. The critical test is whether the agent can propose, test, and revise structural hypotheses about which options share generative factors.

Deep Q-Networks [31] occupy an instructive boundary. Representation learning in hidden layers can implicitly discover shared features as a side effect of gradient descent. However, this occurs within a fixed computational architecture, without an explicit structural learning cycle: the agent does not hypothesize that certain state-action pairs share latent structure, test that hypothesis against held-out data, and revise the computational graph when the hypothesis fails. Under strict DLN definitions, DQN performs high-performance parameter optimization within a fixed topology. Implicit representation learning may place DQN at the Linear-Network boundary, but the absence of explicit structural revision places it in Linear-Plus.

The taxonomy generates a testable prediction: Linear-Plus models should outperform pure Linear when designer-specified structure matches the true generating process, but degrade when structure is absent or changes. Network models should be robust to misspecification because they can revise their structural hypotheses. To test whether DLN stage classifications predict behavior across algorithmically distinct implementations, we simulate two agents per stage using different mechanisms: *ε*-greedy and UCB1 (Linear), Thompson-Factor and RW-Feature (Linear-Plus), HierBayes and LatentFactorEM (Network without DLN mechanisms). All agents were drawn from models with established cognitive science foundations: confidence-based exploration (UCB1; cf. optimism in the face of uncertainty), Bayesian posterior sampling (Thompson Sampling; [13]), associative feature learning (Rescorla-Wagner; [35]), hierarchical Bayesian shrinkage [14], and latent cause discovery [14]. The broader taxonomy (Table 3) extends the classification to machine learning models that occupy the same stages, suggesting the topology distinction is substrate-independent. If agents within the same stage exhibit the same failure/success patterns despite different algorithmic implementations, the classification reflects representational topology rather than algorithmic choice.

## 5 Simulation design

### 5.1 Structural learning cycle

The Network stage is defined as an explicit revision cycle over a small model space (Section 2.4):

1. **Structural hypothesis (Bayesian prior over structure)**. Start with a compressed factor hypothesis for outcome prediction in *G*_*F*_ .
2. **Predictive test (frequentist check)**. Maintain a rolling window of factor prediction errors (mean squared error over the last *W* steps). If the rolling error exceeds an expansion threshold *τ*_expand_, treat this as evidence that the factor hypothesis is inadequate.
3. **Expansion on mismatch**. On mismatch, expand to a tabular representation *G*_tab_ (allocate per-option outcome estimates). This removes compression but restores predictive adequacy when the factor hypothesis is wrong.
4. **Contraction (return transition)**. While operating in the expanded model, periodically test whether a compressed factor model can again predict well. After a holdout period *n*_contract_, open an evaluation window of length *w* in which the agent runs a *shadow* factor model, initialized from current tabular estimates (group means over options that share a factor). If the shadow factor model satisfies MSE_shadow_ *<* (1 − *θ*_contract_)MSE_tab_, the agent contracts back to *G*_*F*_ and (optionally) frees tabular state, returning memory scaling to *O*(*F*).

We report three Network ablations that isolate components of this Level 2 cycle: Network-NoTest disables the predictive check; Network-NoUpdate runs the check but disallows transitions; Network-NoContract allows expansion but disables the return transition, making revision irreversible once the model expands.

### 5.2 Environment

We use a bandit-like environment with *K* options and *F* factors. Each option *i* carries a factor label *c*_*i*_. Outcomes are Gaussian:

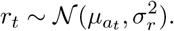

In the structured (prior-correct) condition, 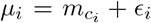 with small idiosyncratic noise; in the unstructured (prior-wrong) condition, the *µ*_*i*_ are independent. Stakes loadings are factor-structured:

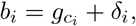

with small idiosyncratic noise. Utility is the sum of outcomes minus incremental stakes penalties.

### 5.3 Agents

Table 4 summarizes the eleven simulated agents, organized by DLN stage. For each non-DLN stage we simulate two algorithmically distinct agents that share identical representational topology, enabling a within-stage consistency test: if agents within the same stage exhibit the same qualitative failure/success pattern under stakes despite different learning algorithms, the DLN classification reflects representational topology rather than algorithmic choice. All baseline agents were selected from models with established cognitive science or behavioral science foundations.

**Table 4:** Simulated agents organized by DLN representational stage. *E*_*t*_: tracks cumulative exposure state. *Cycle*: implements the structural learning cycle (hypothesis → test → update/expand). Network-DLN agents are the only agents possessing both mechanisms.

| DLN Stage | Agent | Mechanism | Memory | $E_t$ | Cycle |
| --- | --- | --- | --- | --- | --- |
| Dot | Dot | Uniform random (no learning) | $O(1)$ | — | — |
| Linear | Linear | ε-greedy, per-option sample means | $O(K)$ | — | — |
| Linear | UCB1 | Upper confidence bounds | $O(K)$ | — | — |
| Linear-Plus | Thompson-Factor | Bayesian posterior sampling, factor posteriors | $O(F)$ | — | — |
| Linear-Plus | RW-Feature | Rescorla-Wagner delta rule, factor weights | $O(F)$ | — | — |
| Network (std) | HierBayes | Hierarchical Normal-Normal, shrinkage | $O(F+K)$ | — | — |
| Network (std) | LatentFactorEM | Online EM, soft factor assignments | $O(KF)$ | — | — |
| Network (DLN) | Network-Full | Factor estimates + expand/contract cycle | $O(F) \rightarrow O(K)$ | ✓ | ✓ |
| Network (DLN) | NoContract | Full, no return transition | $O(F) \rightarrow O(K)$ | ✓ | partial |
| Network (DLN) | NoTest | Full, no predictive check | $O(F)$ | ✓ | — |
| Network (DLN) | NoUpdate | Full, check without transitions | $O(F)$ | ✓ | — |

**Table 5:**
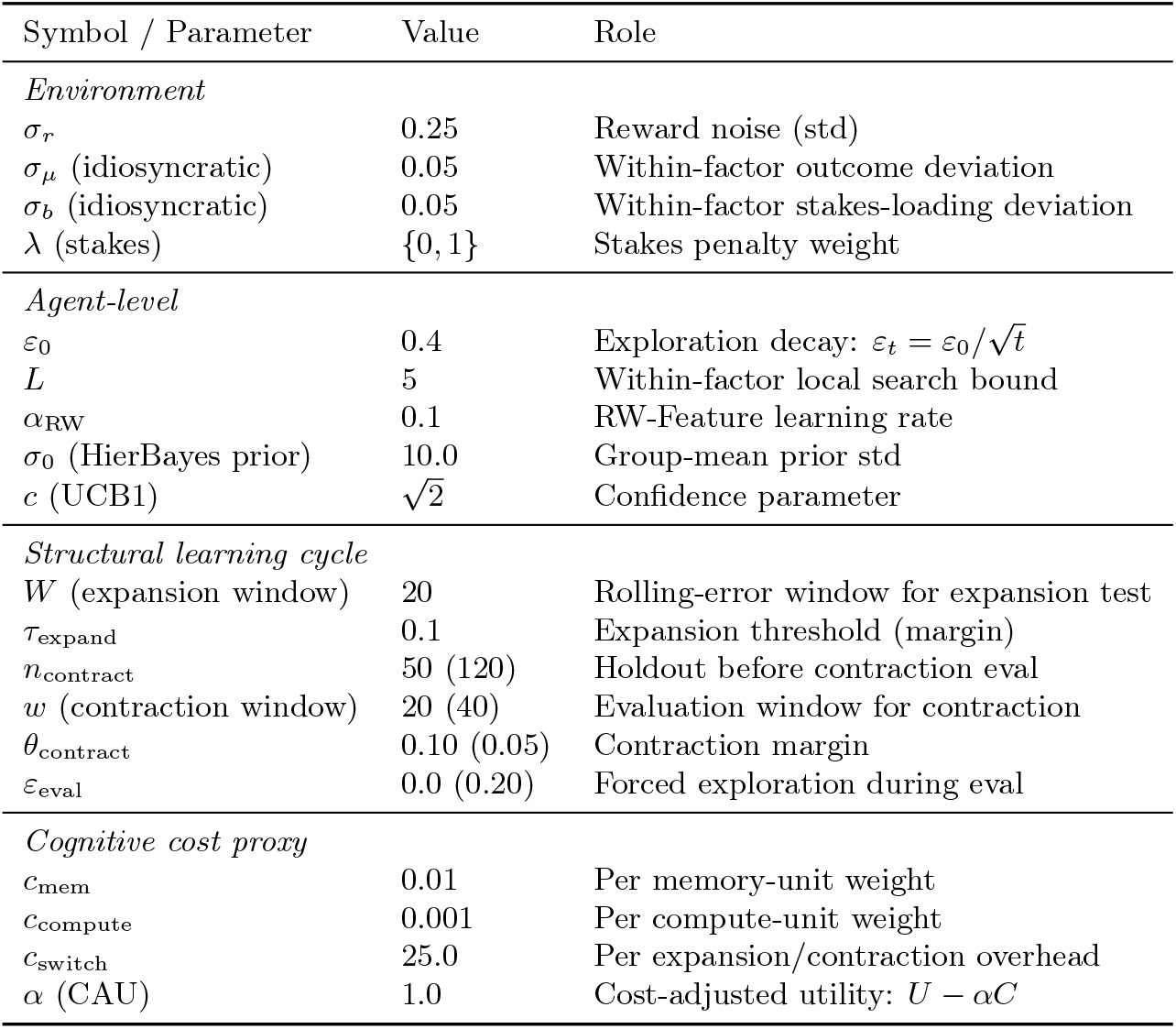
Simulation hyperparameters. All values apply to the paper preset (100 seeds, *K* ∈ {20, 50, 100, 200}, *F* =5, *T* =80); recovery preset differences noted in parentheses.

| Symbol / Parameter | Value | Role |
| --- | --- | --- |
| *Environment* |  |  |
| $\sigma_r$ | 0.25 | Reward noise (std) |
| $\sigma_\mu$ (idiosyncratic) | 0.05 | Within-factor outcome deviation |
| $\sigma_b$ (idiosyncratic) | 0.05 | Within-factor stakes-loading deviation |
| $\lambda$ (stakes) | $\{0, 1\}$ | Stakes penalty weight |
| *Agent-level* |  |  |
| $\varepsilon_0$ | 0.4 | Exploration decay: $\varepsilon_t = \varepsilon_0/\sqrt{t}$ |
| $L$ | 5 | Within-factor local search bound |
| $\alpha_{RW}$ | 0.1 | RW-Feature learning rate |
| $\sigma_0$ (HierBayes prior) | 10.0 | Group-mean prior std |
| $c$ (UCB1) | $\sqrt{2}$ | Confidence parameter |
| *Structural learning cycle* |  |  |
| $W$ (expansion window) | 20 | Rolling-error window for expansion test |
| $\tau_{\text{expand}}$ | 0.1 | Expansion threshold (margin) |
| $n_{\text{contract}}$ | 50 (120) | Holdout before contraction eval |
| $w$ (contraction window) | 20 (40) | Evaluation window for contraction |
| $\theta_{\text{contract}}$ | 0.10 (0.05) | Contraction margin |
| $\varepsilon_{\text{eval}}$ | 0.0 (0.20) | Forced exploration during eval |
| *Cognitive cost proxy* |  |  |
| $c_{\text{mem}}$ | 0.01 | Per memory-unit weight |
| $c_{\text{compute}}$ | 0.001 | Per compute-unit weight |
| $c_{\text{switch}}$ | 25.0 | Per expansion/contraction overhead |
| $\alpha$ (CAU) | 1.0 | Cost-adjusted utility: $U - \alpha C$ |

#### Dot stage

The Dot agent selects uniformly at random and performs no learning. It serves as the null-graph baseline: no belief state persists across encounters, and neither option-factor nor stakes-factor structure is represented.

#### Linear stage (Linear, UCB1)

Both agents maintain *K* independent option-level statistics with no cross-option information sharing—the defining property of Linear-stage topology (Section 2.1). Linear employs *ε*-greedy selection over frequentist sample means; action selection scores each option using a local stakes heuristic that ignores the cross-term 2*E*_*t*_*b*_*i*_:

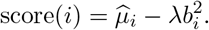

This scoring function treats hedging as separable across options and does not track cumulative exposure. A hybrid agent that tracks *E*_*t*_ while maintaining independent option beliefs was considered but excluded: tracking cumulative exposure requires representing how past choices affect future evaluations, a cross-option temporal dependency inconsistent with the branch-by-branch independence that defines Linear-stage cognition. Isolating the factor-compression and exposure-tracking contributions via such a baseline is a direction for future work. UCB1 employs deterministic selection via upper confidence bounds 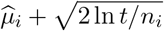, implementing the principle of optimism in the face of uncertainty—a mechanism that has received substantial attention as a computational account of human directed exploration [13]. The two agents differ algorithmically (undirected versus directed exploration) but share identical representational topology: a null belief graph on *K* option nodes in which evidence about option *i* cannot update beliefs about option *j*.

#### Linear-Plus stage (Thompson-Factor, RW-Feature)

The Linear-Plus category captures an empirically important intermediate between Linear and Network: agents that share information across options through designer-specified structure (here, factor labels *c*_*i*_) but neither discover that structure from data nor revise it in response to predictive failure. This distinguishes Linear-Plus from Linear, where no cross-option sharing occurs, and from Network, where the agent can propose, test, and revise structural hypotheses. The critical boundary is structural revision: Linear-Plus agents cannot detect that their factor hypothesis is inadequate, cannot expand to option-level estimation when it fails, and cannot contract back when structure returns.

Thompson-Factor implements Bayesian posterior sampling over factor-level parameters via Normal–Normal conjugate updates, selecting actions by Thompson sampling (draw from each factor posterior, select the best factor, then local search over at most *L*=5 options). Thompson sampling is a standard algorithm in computational accounts of human exploratory behavior [13, 37]. RW-Feature implements the Rescorla–Wagner delta rule [35]—one of the most extensively validated models in associative learning and behavioral neuroscience—applied here to factor-level feature weights 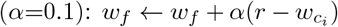. Both agents exploit given factor structure to achieve *O*(*F*) memory scaling, use the same local 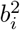 stakes heuristic as Linear, and neither tracks cumulative exposure *E*_*t*_ nor monitors structural adequacy.

#### Network, standard (HierBayes, LatentFactorEM)

Both agents implement forms of cross-option structure learning that have been studied extensively in computational neuroscience and cognitive science, but lack the DLN-specific mechanisms of cumulative exposure tracking and the structural learning cycle.

HierBayes implements a hierarchical Normal–Normal model in which per-factor posteriors are shrunk toward a learned group mean (*µ*_0_=0, *σ*_0_=10, *τ* =1), with action selection via Thompson sampling from shrinkage-adjusted posteriors and local *L*=5 search. Hierarchical Bayesian shrinkage is central to computational accounts of human generalization from limited data [14, 44]: sharing statistical strength across related items enables rapid inference that neither purely local nor purely global estimation can achieve.

LatentFactorEM infers soft factor assignments via online Expectation-Maximization with a log-space E-step for numerical stability, instantiating the latent cause inference framework that has been proposed to explain human category learning and context-dependent memory retrieval [14].

Both agents discover or refine latent structure, placing them in the Network stage. However, neither tracks *E*_*t*_ nor implements the hypothesis → test → update/expand cycle that defines DLN Network cognition. This pairing tests whether hierarchical structure learning alone—without the variable-compression channel and structural revision mechanism—is sufficient under stakes and regime change.

#### Network, DLN (Network-Full and ablations)

The Network-Full agent maintains factor-level outcome estimates 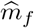 and factor-level exposure estimates 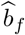, tracks *E*_*t*_, and uses the true marginal penalty form. Within a chosen factor, Network searches over at most *L*=5 options to avoid hiding *O*(*K*) scans in factor selection. It also runs the predictive check, can expand to tabular outcome estimates when the factor hypothesis fails, and can contract back to the factor model when predictive evidence supports compression again (completing a return transition in the revision graph).

Three ablations isolate components of the Level 2 cycle: Network-NoTest disables the predictive check; Network-NoUpdate runs the check but disallows model-class transitions; Network-NoContract allows expansion but disables the return transition to the compressed model.

### 5.4 Metrics and statistics

Per episode we report: (i) total outcome, (ii) total stakes penalty, (iii) utility *U* = ∑_*t*_(*r*_*t*_ − Δ*P*_*t*_), (iv) cognitive cost proxy *C*, (v) cost-adjusted utility *U* − *αC*, (vi) jointly efficient action pick count, (vii) number of expansion transitions *G*_*F*_ → *G*_tab_, (viii) number of contraction (return) transitions *G*_tab_ → *G*_*F*_, and (ix) time spent in each model class (steps in *G*_*F*_ vs. *G*_tab_).

Results are averaged over 100 random seeds with paired comparisons across seeds (same environment seed per agent). For key paired differences we report bootstrap 95% confidence intervals (percentile bootstrap over seeds) and paired Cohen’s *d* on the seed-wise difference.

## 6 Results

### 6.1 Option-factor structure compression (Claim C1)

Table 6 reports cost-adjusted utility in the stable structured environment (stakes off; Figure 2). As *K* grows, Linear cognitive cost scales with *K*, while Network cost is approximately constant in *F* (Table 7). Network overtakes Linear for larger option sets: at *K* = 200, Network-Full achieves cost-adjusted utility 80.47 vs. Linear 64.62 (paired difference mean +15.85).

**Table 6:** Cost-adjusted utility (mean ± sd) in the stable structured environment (stakes=0).

| $K$ | Dot | Linear | Network-Full |
| --- | --- | --- | --- |
| 20 | $0.96 \pm 8.57$ | $89.61 \pm 35.14$ | $89.17 \pm 41.44$ |
| 50 | $0.72 \pm 9.00$ | $82.44 \pm 35.37$ | $88.25 \pm 29.95$ |
| 100 | $1.58 \pm 8.34$ | $73.28 \pm 34.09$ | $89.18 \pm 31.66$ |
| 200 | $-0.94 \pm 9.08$ | $64.62 \pm 34.45$ | $80.47 \pm 28.52$ |

**Table 7:**
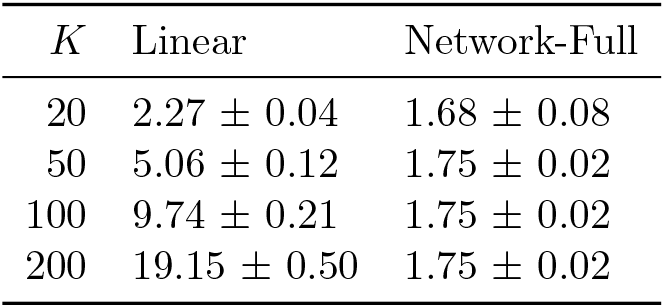
Cognitive cost proxy (mean ± sd) showing *O*(*K*) scaling for Linear and *O*(*F*) scaling for Network.

**Figure 1.**
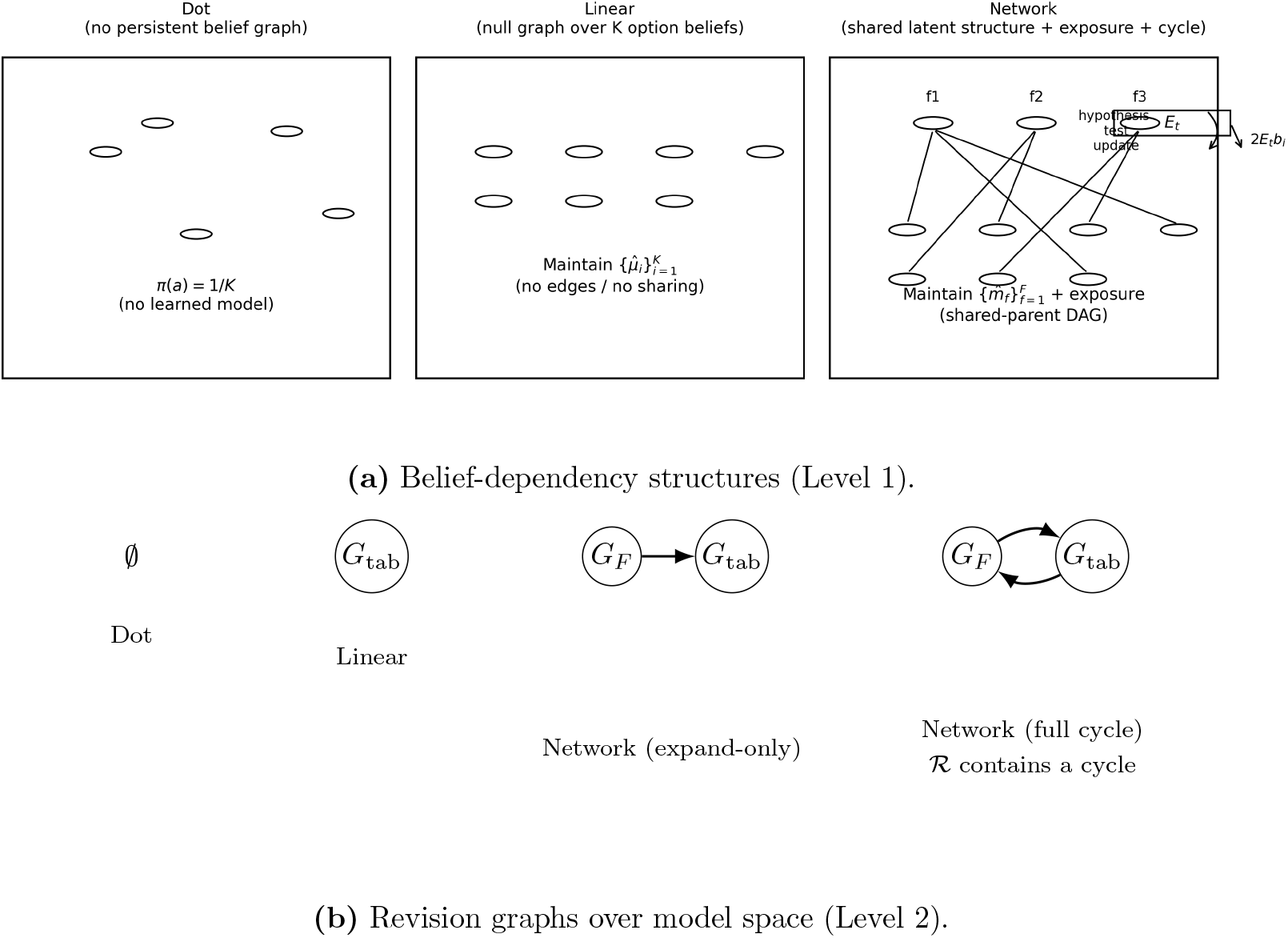
(a) Belief-dependency structures (Level 1). Dot: no persistent graph. Linear: null graph on *K* option nodes. Network: bipartite factor DAG with *F* factor nodes connected to *K* options. (b) Revision graphs over model space (Level 2). Dot: empty model space. Linear: fixed point in model space (no revision edges). Network (expand-only): directed path *G*_*F*_ → *G*_tab_. Network (full cycle): two directed transitions (expand and contract) forming a cycle in ℛ, enabling bounded recovery (Proposition 1(iii)).

**Figure 2.**
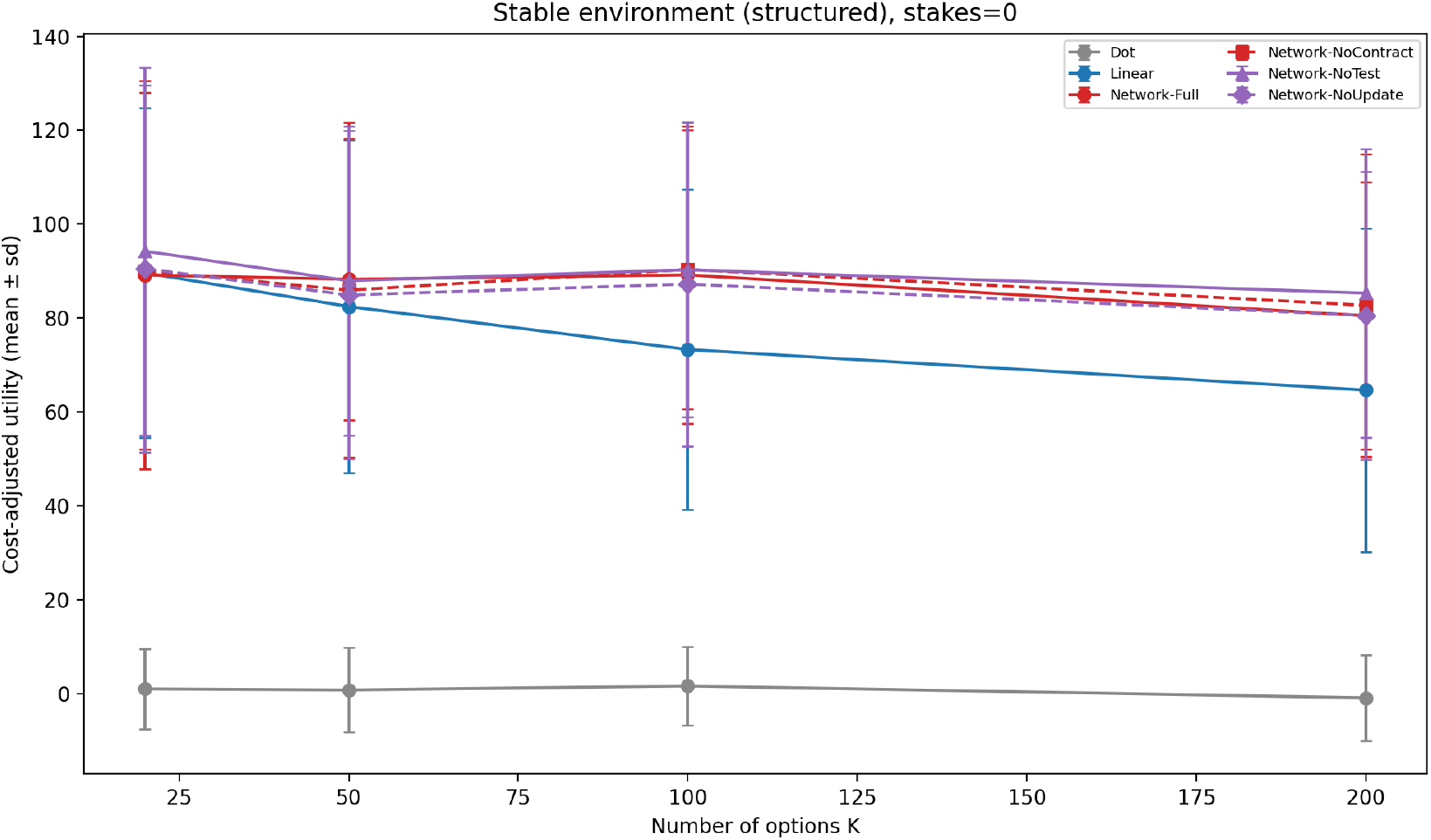
Stable structured environment (stakes=0): cost-adjusted utility vs. number of options *K*. Network overtakes Linear at larger *K* due to *O*(*F*) cost scaling.

#### Empirical vs. analytic crossover

Proposition 1(i) predicts a crossover at *K*^∗^ = *F* +*c*_meta_/*c*_param_ = 5 + 25.0/0.01 = 2,505. The empirical crossover—the *K* at which Network CAU first exceeds Linear CAU—occurs near *K* ≈ 22 (by linear interpolation between the *K*=20 and *K*=50 grid points), roughly 113× earlier. The gap arises because the analytic formula captures only the *cost* component of the advantage (*O*(*F*) vs. *O*(*K*) memory), whereas factor pooling also improves *prediction quality* —a statistical shrinkage effect analogous to James–Stein estimation [20]. Decomposing the Network–Linear advantage into utility and cost components confirms that the utility gain (better reward estimates from factor pooling) dominates the total advantage at moderate *K*, while the cost saving becomes the primary driver only at large *K*. The analytic *K*^∗^ therefore provides an upper bound on the crossover: the actual advantage emerges much earlier because compression improves both computation *and* estimation.

### 6.2 Stakes-factor structure compression (Claim C2)

With stakes active, Linear agents incur large penalties because they do not represent the cross-term 2*E*_*t*_*b*_*i*_ and cannot hedge cumulative exposure. Network agents track *E*_*t*_ and learn factor-level exposure structure, enabling hedging and keeping penalty bounded. At *K* = 200 in the structured setting, Network-Full maintains positive mean utility (69.50) while Linear incurs substantial negative utility (−706.90). The paired utility difference is +776.40.

### 6.3 Dot baseline

Dot serves as the null-graph baseline: it does not accumulate value estimates, does not represent option-factor or stakes-factor structure, and does not track cumulative exposure *E*_*t*_. As a result, Dot does not benefit from increasing *K* even when option-factor structure is present. In the stable structured setting with stakes off (Table 6), Dot’s cost-adjusted utility remains near zero across *K* (means between −0.94 and 1.58), reflecting near-chance outcomes combined with near-zero cognitive cost.

With stakes active (Table 8), Dot incurs negative utility (roughly −81 to −129) because cumulative exposure follows an uncontrolled random walk. Dot avoids Linear’s extreme degradation only because it also avoids systematic exploitation; this reflects robustness through non-commitment rather than compression. Network differs by maintaining positive utility while simultaneously optimizing outcomes and controlling marginal exposure impact.

**Table 8:** Utility (mean ± sd) in the structured environment with stakes (*λ*=1), organized by DLN stage. All non-DLN agents collapse to negative utility; the collapse is worst for Linear-Plus agents, which aggressively exploit factor structure without tracking cumulative exposure *E*_*t*_. Only Network-DLN maintains positive utility.

| DLN Stage | Agent | $K=20$ | $K=50$ | $K=100$ | $K=200$ |
| --- | --- | --- | --- | --- | --- |
| Dot | Dot | $-129 \pm 189$ | $-81 \pm 102$ | $-82 \pm 125$ | $-102 \pm 160$ |
| Linear | Linear | $-1,192 \pm 1,565$ | $-1,070 \pm 1,737$ | $-897 \pm 1,439$ | $-707 \pm 1,267$ |
| Linear | UCB1 | $-425 \pm 564$ | $-83 \pm 147$ | $-18 \pm 29$ | $-59 \pm 84$ |
| Linear-Plus | Thompson-Factor | $-6,725 \pm 10,661$ | $-5,040 \pm 6,638$ | $-4,352 \pm 5,024$ | $-3,220 \pm 4,159$ |
| Linear-Plus | RW-Feature | $-5,909 \pm 7,858$ | $-4,490 \pm 5,728$ | $-4,618 \pm 5,524$ | $-3,714 \pm 4,557$ |
| Network (std) | HierBayes | $-3,863 \pm 9,209$ | $-2,414 \pm 3,888$ | $-2,639 \pm 3,351$ | $-2,387 \pm 2,839$ |
| Network (std) | LatentFactorEM | $-1,662 \pm 1,972$ | $-1,807 \pm 2,210$ | $-1,697 \pm 1,997$ | $-1,962 \pm 2,455$ |
| Network (DLN) | Network-Full | $69 \pm 37$ | $74 \pm 37$ | $71 \pm 33$ | $70 \pm 28$ |

### 6.4 Linear-Plus agents: factor compression without DLN mechanisms

The two Linear-Plus agents—Thompson-Factor and RW-Feature—share the same factor-level representation as Network but lack the DLN structural cycle and cumulative exposure tracking (Table 4). Their results isolate whether factor compression alone accounts for the Network advantage, or whether the variable-compression channel (*E*_*t*_ tracking) and the learning cycle are essential.

#### Stable environment with stakes (*λ*=1)

Both Linear-Plus agents collapse far worse than Linear (Figure 3): at *K*=200, Thompson-Factor reaches CAU = −3,222 and RW-Feature reaches −3,715, compared to Linear at − 726 and Network-Full at +68. The failure mechanism is shared: both agents use a local 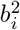 heuristic and do not track cumulative exposure *E*_*t*_. Without the cross-term 2*E*_*t*_*b*_*i*_, they cannot hedge—and their factor-level exploitation *accelerates* exposure accumulation by concentrating choices in high-reward factors. This confirms that the Network advantage under stakes (Claim C2) arises specifically from exposure tracking, not from factor compression alone.

**Figure 3.**
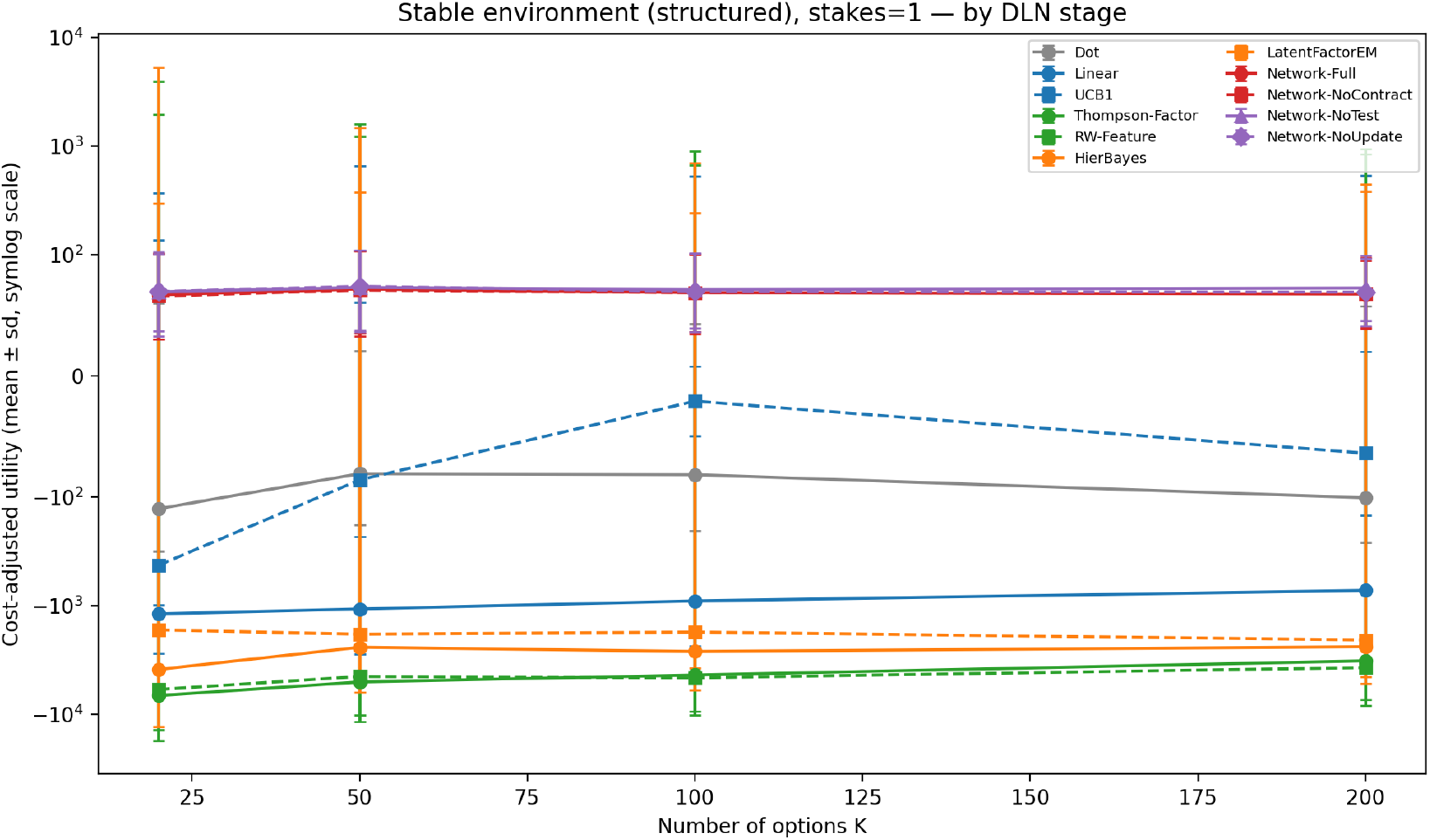
Cost-adjusted utility vs. *K* in the structured stakes condition (*λ*=1, symlog scale). Agents grouped by DLN stage (Table 4). All non-DLN agents collapse to negative CAU; Linear-Plus (green) collapses worst due to factor-level exploitation without *E*_*t*_ tracking. Only Network-DLN (red) maintains positive CAU.

#### Regime change with stakes (*λ*=1)

Under the recovery condition (regime change at step 100), the failure intensifies by an order of magnitude: Thompson-Factor reaches CAU ≈ −49,000 and RW-Feature reaches ≈ −38,000 (Figure 6, center panel), compared to +60 to +89 for Network-Full and −9,400 to −11,300 for Linear (at *K*=200). Neither Linear-Plus agent possesses the expansion mechanism that allows Network to detect model failure and revert to tabular estimates; both continue to exploit stale factor representations learned during the unstructured phase, compounding both prediction error and exposure penalties.

#### Interpretation

Factor-level compression is necessary but not sufficient. Agents that share the same factor representation as Network but lack DLN mechanisms perform *worse* than even the option-level Linear agent under stakes, and catastrophically worse under regime change. The DLN contribution beyond shared factor compression is twofold: (i) the variable-compression channel, representing how past choices affect future evaluations via the cross-term 2*E*_*t*_*b*_*i*_; and (ii) the structural learning cycle, enabling model-class transitions when the environment invalidates the current representation. That two algorithmically distinct Linear-Plus agents (Bayesian posterior sampling and Rescorla–Wagner associative learning) exhibit the same failure pattern confirms that the deficit is structural, not algorithmic.

### 6.5 Within-stage consistency

The DLN taxonomy predicts that agents classified in the same stage should exhibit the same qualitative failure/success pattern under stakes, regardless of their learning algorithm. To test this prediction, we simulate two algorithmically distinct agents per non-DLN stage (Table 4) and compare their behavior in the structured environment with stakes (*λ*=1).

Under stakes, the pattern is consistent within each stage and sharply differentiated across stages (Figure 3). Both Linear-stage agents (Linear, UCB1) collapse to negative cost-adjusted utility as cumulative exposure penalties dominate. Both Linear-Plus agents (Thompson-Factor, RW-Feature) collapse further—aggressive factor-level exploitation without *E*_*t*_ tracking accelerates exposure accumulation, amplifying the penalty cross-term. Both Network-standard agents (HierBayes, LatentFactorEM) collapse as well, despite implementing hierarchical structure learning: without the variable-compression channel, they cannot hedge cumulative exposure. Only Network-DLN agents maintain positive cost-adjusted utility across all *K*.

Under regime change with stakes (Figure 6, center panel), the stage ordering sharpens. Linear-Plus agents (−38,000 to −49,000 at *K*=200) collapse worse than Network-standard agents (−21,000 to −28,000), which collapse worse than Linear agents (−1,000 to −9,000), which collapse worse than Dot (−316). This inverted ordering—more sophisticated factor exploitation producing worse outcomes—is a mechanistic prediction of the DLN framework: factor-level exploitation concentrates choices in high-reward factors, which accelerates cumulative exposure accumulation in the absence of marginal impact tracking. Network-DLN agents (+60 to +106) are the sole survivors.

These results support two conclusions. First, the DLN stage classification captures a structural property—representational topology—that is invariant across learning algorithms: the failure mode is determined by what an agent represents, not how it learns within that representation. Second, the absence of *E*_*t*_ tracking is the critical deficit shared by all non-DLN agents, confirming that the variable-compression channel is the mechanism that separates Network-DLN from all other stages under stakes.

### 6.6 Jointly efficient actions (Claim C3)

In the structured stakes condition, the environment contains a sparse set of actions that are *jointly efficient* by construction: they are high on expected outcome *and* reduce marginal exposure impact under the stakes objective. Formally, the true incremental penalty for selecting option *i* at exposure state *E*_*t*_ is

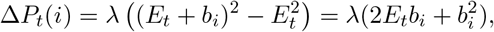

so the *marginal* stakes contribution depends on the sign interaction between the cumulative exposure *E*_*t*_ and the option’s loading *b*_*i*_. A jointly efficient action is one that simultaneously has high *µ*_*i*_ and (when *E*_*t*_ is positive) has *b*_*i*_ *<* 0 so that the cross-term 2*E*_*t*_*b*_*i*_ is negative.

#### Evaluation definition

At environment initialization we flag a small subset of options as jointly efficient (the `dual_mask`): options in the top reward quantile that also have negative exposure loading. This flag is computed from the environment’s ground-truth (*µ*_*i*_, *b*_*i*_) and is fixed *before* any agent acts. We report the pick rate as the fraction of timesteps in which the chosen action lies in this flagged set (Figure 4).

**Figure 4.**
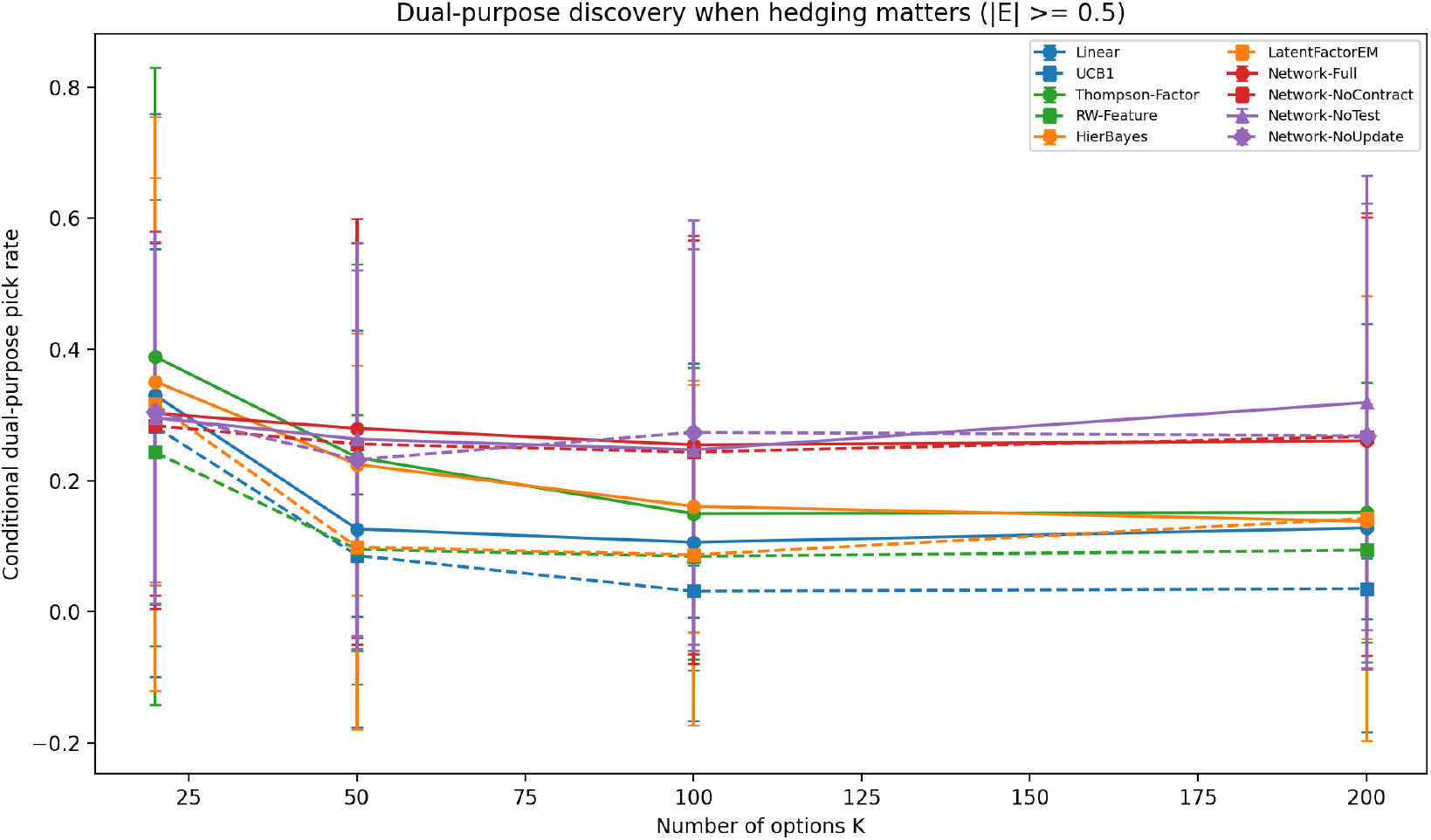
Jointly efficient action pick rate (structured, stakes=1).

#### Why intersection requires representing both decompositions

To identify a jointly efficient action, a policy must (i) know which actions are promising on expected outcomes (Option-factor structure) and (ii) evaluate *marginal* stakes impact given current exposure *E*_*t*_ (Stakes-factor structure). Network-Full represents both: it learns factor-level outcome estimates, learns factor-level exposure estimates, tracks *E*_*t*_, and scores factors/actions using the true marginal form. Linear fails on (ii): it uses a local stakes heuristic 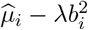 that ignores the cross-term and the sign interaction with *E*_*t*_, so it systematically undervalues actions that are beneficial precisely because their exposure loading offsets existing exposure. Dot fails on both (i) and (ii) because it is non-learning and exposure-agnostic.

### 6.7 Prior-wrong condition: the learning cycle expands the model

In the unstructured condition, the factor hypothesis for outcome prediction is incorrect. Network-Full allocates tabular state (expansion) and improves raw utility relative to NoTest/NoUpdate ablations, while paying an explicit cognitive cost for doing so. For example at *K* = 200 (unstructured, stakes off), Network-Full exceeds Network-NoTest by +5.04 utility (bootstrap 95% CI [0.89, 9.42]) and exceeds Network-NoUpdate by +5.05 (bootstrap 95% CI [0.59, 9.66]), while exhibiting more hypothesis expansions (Figure 5). Because expansion incurs a one-time *O*(*K*) cognitive cost, the net cost-adjusted advantage depends on the time horizon: at *T* =80 the fixed cost dominates, but in the longer regime-change episodes (*T* =300; Section 6.8) the accumulated reward gain amortizes the expansion cost and Network-Full surpasses all ablations on CAU.

**Figure 5.**
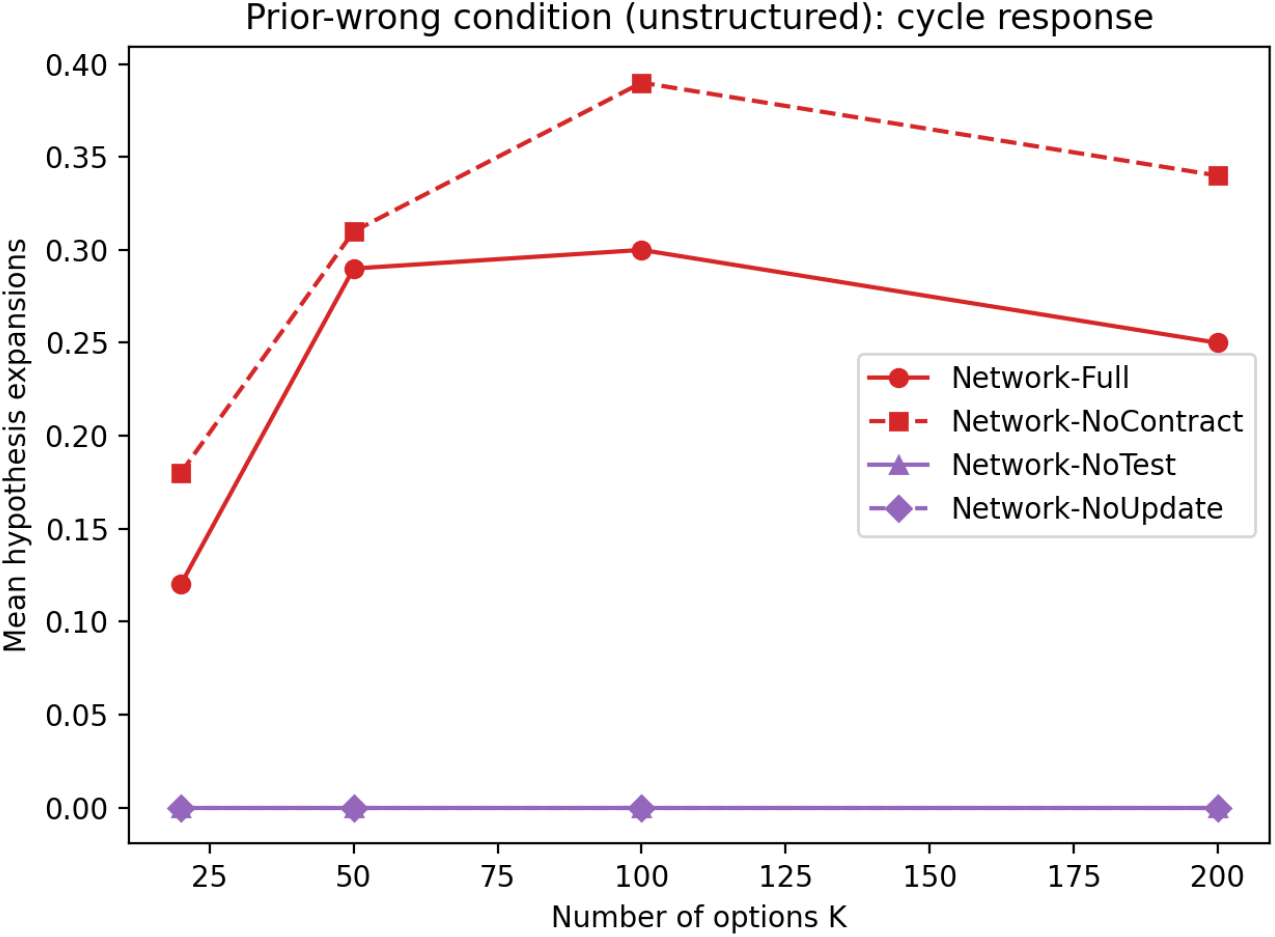
Prior-wrong condition (unstructured, stakes=0): mean number of hypothesis expansions (switches) for Network variants.

**Figure 6.**
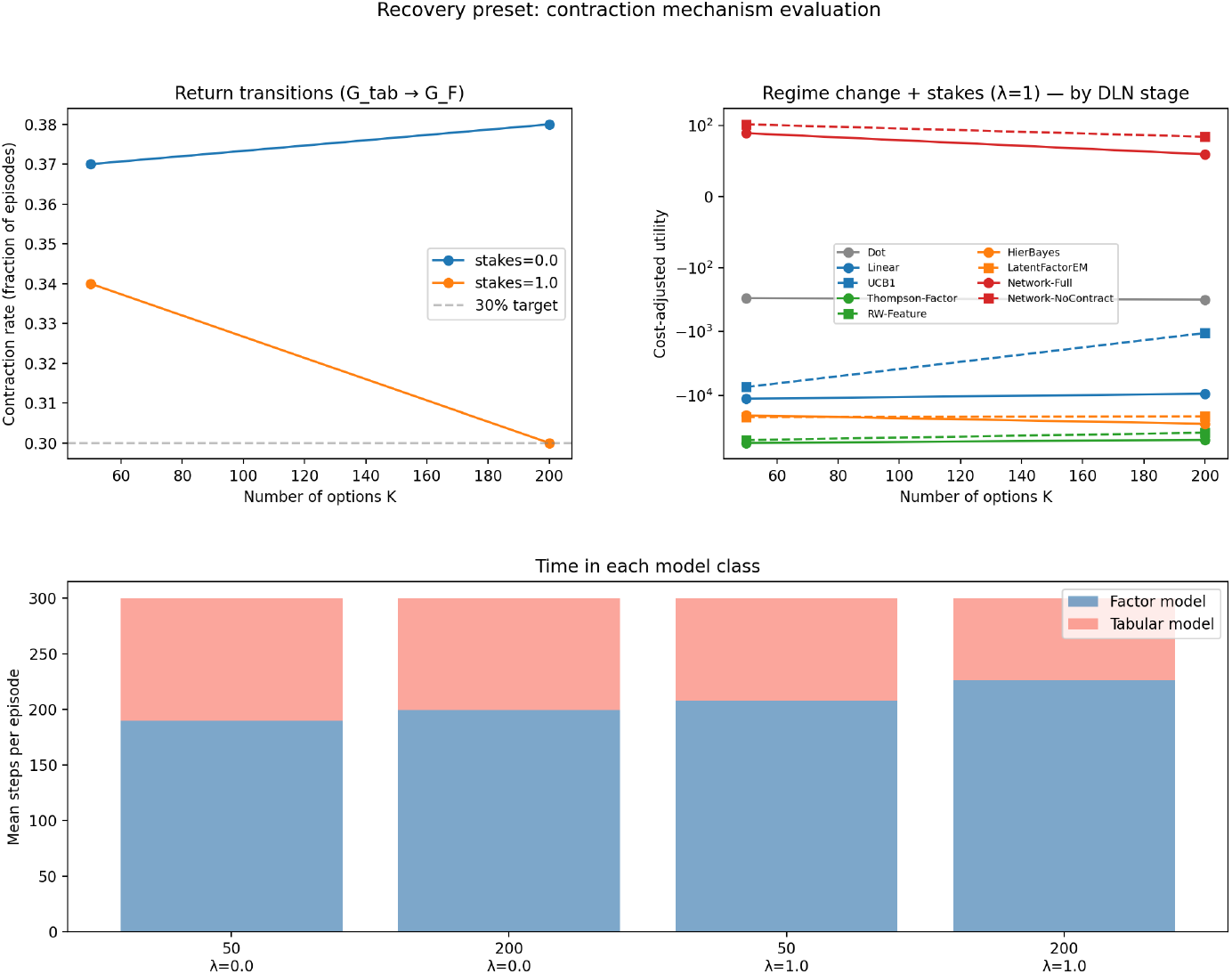
Recovery condition (regime change at step 100). *Top left:* Contraction rate vs. *K*; dashed line marks 30% target. *Top right:* Cost-adjusted utility (*λ*=1, symlog scale) by DLN stage (Table 4). Stage ordering inverts: Linear-Plus collapses worst (∼−4×10^4^), then Network-standard, Linear, Dot. Only Network-DLN maintains positive CAU. *Bottom:* Time allocation in factor vs. tabular model.

#### Interpreting the ablations

Network-NoTest corresponds to compression without validation: it trusts the factor hypothesis unconditionally and therefore continues pooling across irrelevant labels when the prior is incorrect, accumulating systematic bias. Network-NoUpdate corresponds to validation without adaptation: it allocates computation (and may even allocate tabular state for comparison) to detect mismatch, but is not permitted to switch its action model, so it pays overhead without recovering performance. Network-Full requires both the predictive test and the update/expansion branch to recover under prior failure; the performance gaps in the unstructured condition demonstrate that the *learning cycle* (not just factorization) is performing work.

### 6.8 Contraction and bounded recovery (return transition)

The prior-wrong condition above isolates why the Network stage must include *expansion*: when the factor hypothesis fails, an agent that remains in *G*_*F*_ is locked into a mis-specified compressed representation. Completing the revision cycle requires an additional capability: once compression becomes appropriate again, the agent must be able to return to *G*_*F*_ .

To isolate this return transition, we include Network-NoContract, which is identical to Network-Full except that it disables contraction (*G*_tab_ → *G*_*F*_). In the static conditions above (Sections 6.1–6.7), contraction does not activate: structure is either always present (no expansion) or always absent (no opportunity to contract). Network-Full and Network-NoContract therefore exhibit equivalent performance in those experiments.

#### Regime-change condition

To exercise the contraction mechanism, we introduce a *recovery* condition: a regime-change environment with *T* =300 in which rewards are initially unstructured (steps 1–99) and switch to a structured factor model at step 100. This forces expansion during the unstructured phase and creates a post-change window (200 steps) sufficient for the contraction holdout (*n*_contract_=50) and evaluation window (*w*=20).

Network-Full correctly detects the wrong hypothesis and expands to *G*_tab_ in 36–53% of episodes (depending on condition). After structure onset, the contraction mechanism opens 0.36–0.53 evaluation windows per episode, confirming that the holdout-then-evaluate cycle operates as designed. With a wider evaluation window (*w*=40), a lower contraction margin (*θ*_contract_=0.05), fresh shadow-model initialization (not seeded from contaminated tabular estimates), and modest forced exploration during evaluation (*ε*_eval_=0.20), successful contractions occur in 30–38% of episodes across conditions (Figure 6, left panel). Three implementation corrections were necessary to reach this rate: (i) the shadow model’s prediction error was previously computed using *post-update* Q-values, yielding artificially near-zero MSE for the tabular model; (ii) on-policy evaluation only tested options the tabular model already calibrated well, masking the factor model’s advantage on unvisited options; and (iii) initializing the shadow model from pre-regime-change tabular estimates contaminated its priors.

The evaluation mechanism carries a non-trivial cost: forced exploration during the holdout window reduces immediate reward. At *K*=200 with stakes (*λ*=1), Network-NoContract achieves CAU = 84.3 compared to 59.7 for Network-Full—a −24.7 difference attributable to evaluation overhead. Both agents vastly outperform Linear, which collapses under the regime change. Without stakes (*λ*=0), the pattern is similar (Network-NoContract 129.6 vs. Network-Full 111.3).

These results provide support for Proposition 1(iii) at the *architectural* level: the contraction mechanism correctly identifies when the factor model outperforms the tabular model and executes the return transition at meaningful rates. The net CAU impact is currently negative because the evaluation cost (forced exploration) exceeds the recovery benefit within the 200-step post-change runway. Longer episodes, lower exploration rates, or amortization across multiple regime changes would shift this tradeoff. The key contribution is demonstrating that model-class reversion—a Level 2 operation in ℛ—is feasible and fires at empirically meaningful rates.

## 7 Discussion

### 7.1 Interpretation

Under Assumptions A1–A6, the simulation isolates the efficiency consequences of representing shared structure. Dot is included as the null-graph baseline (no accumulation, no compression) to anchor the three-stage ordering; it is expected to ignore both structure and stakes. The revision graph ℛ introduced in Section 2.4 provides a formalization of this deployment layer: the structural learning cycle implements Level 2 computation (navigation of ℛ), and the crossover condition *K*^∗^ = *F* + *c*_meta_/*c*_param_ determines when investing in Level 2 is cost-effective. This connects to metacognitive monitoring [11, 5] by formalizing what is monitored (model adequacy, via predictive tests) and what actions are available in response (transitions in ℛ). The mapping should be interpreted as follows: *if* DLN Network is interpreted as a topology that supports factor reuse and cross-branch interaction, *then* it should produce *O*(*F*)-like scaling advantages and the ability to identify jointly efficient actions.

#### Meta-cognition as a property of model space

DLN stages differ not only in the structure of the belief graph *G* (Level 1) but in whether the agent can navigate a non-trivial model space (Level 2). Linear cognition is a fixed point in ℳ. Network cognition traverses ℳ via revision transitions, including a return transition whose functional consequence is bounded cost recovery after model failure (Proposition 1(iii)). This provides a foundation for instantiating DLN in other domains: any system where an observer, agent, or learner can be characterized by (i) a current representational structure and (ii) a set of available transitions between structures may be analyzed by the structure of its revision graph.

The Network advantage obtains in stable environments without requiring distribution shift or novelty-seeking [13, 37]; the focus is representational reuse under stationarity.

#### Dissociating compression, exposure tracking, and structural revision

The Linear-Plus and Network-standard agents provide a clean decomposition of the DLN Network advantage across three components. In stable environments without stakes, Linear-Plus agents (Thompson-Factor, RW-Feature) and Network-standard agents (HierBayes, LatentFactorEM) perform comparably to or better than Network-Full because factor-level compression and efficient exploration are orthogonal to the DLN-specific mechanisms. Under stakes, cumulative exposure tracking reverses the ordering by orders of magnitude: all non-DLN agents collapse, while Network-Full maintains positive CAU. Under regime change with stakes, the gap widens further because non-DLN agents cannot detect that their structural hypothesis is wrong. The DLN contribution thus has three components: (i) factor compression (shared by Linear-Plus and Network-standard agents), (ii) cumulative exposure tracking (which only Network-DLN implements), and (iii) structural revision via the learning cycle (which only Network-DLN implements). Non-DLN agents achieve (i) but not (ii) or (iii).

### 7.2 Boundary conditions

The results depend on explicit structural gaps between *K* and *F* and on the presence of cross-terms in the stakes objective.

#### When *K* = *F* (no compression available)

If every option is its own factor, then there is no shared component to compress. In that case, a Network factor representation should offer no advantage over Linear, and may perform worse due to representational overhead. This constitutes a key falsifier for the compression thesis.

#### Partial structure

If option-factor structure is weak (large within-factor deviations), the benefit of shrinkage and factor scoring should degrade smoothly. Likewise for stakes-factor structure: if exposures are not aligned with factors, the hedging benefit should degrade. A structure-strength sweep makes these boundary conditions visible.

#### Stakes scaling versus stakes structure

This paper models stakes through *structured exposures* with a covariance-inducing cross-term. Scaling stakes magnitude alone is a different question; both can be studied, but they test different mechanisms.

### 7.3 Limitations and future work

#### Observed factor labels (A1)

We assume the Network agent has access to a shared-component representation *c*_*i*_. This provides a representational advantage rather than requiring the agent to learn it. *Future direction:* replace *c*_*i*_ with learned clustering or learned latent factors (Bayesian nonparametrics or representation learning) and report the additional sample and cognitive costs required to learn the representation.

#### 1D exposure state (A5)

We use a single exposure dimension to make the cross-term explicit and interpretable. *Future direction:* extend to *d*-dimensional exposures with *E*_*t*_ ∈ ℝ^*d*^ and penalty ∥*E*_*t*_∥^2^ (or a quadratic form 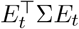). Measure how Network cost scales with the number of exposure factors rather than with the number of options.

#### Cognitive cost proxy (A6)

We use an algorithmic proxy (memory and compute scaling). *Future direction:* perform sensitivity analysis over cost weights and *α*, and report Pareto frontiers over (utility, cost) rather than a single calibrated point estimate.

#### Local search within factor

Network uses a bounded local search of size *L* within a chosen factor to avoid hiding *O*(*K*) scans. *Future direction:* vary *L* (ablation) to quantify the tradeoff between factor-level compression and within-factor deviation exploitation.

#### Developmental scope

Developmental interpretations of DLN stage differences are potentially culturally contingent [18]. The present simulation tests a computational mapping under explicit assumptions; extending the framework to developmental claims requires behavioral validation across populations.

#### Decomposing compression and exposure tracking

The present design confounds two mechanisms: factor-level compression (representing shared structure once) and cumulative exposure tracking (using *E*_*t*_ to compute marginal impact). A Linear agent augmented with *E*_*t*_-tracking but no factor structure would isolate these contributions. *Future direction:* implement a “Linear+*E*_*t*_” baseline and measure how much of the stakes-condition advantage (Claim C2) arises from compression versus exposure tracking.

## 8 Reproducibility

All tables in this paper are derived from outputs/paper/results/episode metrics.csv. The default run that reproduces the CSV and baseline figures is:

`python src/dln_core_variable_cycle.py --preset paper --out outputs/paper`

The run writes `outputs/paper/results/manifest.json` containing hyperparameters, seeds, and file hashes. Baseline figures are written under `outputs/paper/artifacts/figures/`. To regenerate figures with different styling, use the CSV as the single source of truth and rebuild plots externally.

### RNG architecture

The simulation uses `numpy.random.SeedSequence.spawn` to create independent per-agent random streams, so that adding or removing agents does not shift results for other agents. The canonical results correspond to commit `283214e`; all tables and in-text statistics in this manuscript are derived from the committed CSV at that revision.

### Code availability

The complete source code, simulation framework, and reproducibility scripts are available at: https://github.com/aliawu08/dln-compression-model

## 9 Conclusion

Two formal objects carry the DLN distinction: the belief-dependency graph *G*, whose structure determines within-episode inference cost, and the revision graph ℛ over model space, whose structure determines meta-cognitive capacity to revise the active belief structure. Under explicit assumptions, Network policies outperform Linear policies in stable environments once option sets are sufficiently large (compression of option-factor structure), avoid exposure degradation under stakes through factor-level exposure learning and cumulative exposure tracking (compression of stakes-factor structure), and revise their structural hypothesis via expansion and contraction transitions when evidence warrants (learning cycle). Dot remains a baseline regime with no accumulation: it neither benefits from option-factor structure nor controls cumulative exposure, and therefore stays near baseline utility. Proposition 1 derives the crossover *K*^∗^ = *F* + *c*_meta_/*c*_param_ from the interaction between within-episode compression and model-revision overhead, and shows that including a return transition in ℛ provides bounded recovery after model failure. The mapping is falsifiable at the level of assumptions and boundary tests: when shared structure vanishes (*K*=*F*) or when structure is weak, the Network advantage should disappear. Within-stage consistency results (Section 6.5) provide further support for the taxonomy: two algorithmically distinct agents per stage exhibit the same qualitative failure/success pattern under stakes, and the collapse ordering inverts relative to algorithmic sophistication—Linear-Plus agents collapse worst, followed by Network-standard, then Linear—confirming that the classification reflects representational topology rather than algorithmic choice. This two-object characterization—*G* for within-episode inference, ℛ for cross-episode revision—provides a concrete bridge between qualitative DLN descriptions and measurable computational predictions, including in settings beyond decision-making where representational constraints determine what information an agent can extract from its environment.

